# Competition in the savanna: models of species assemblages in Kruger National Park, South Africa

**DOI:** 10.1101/147652

**Authors:** Sadie J. Ryan, Joshua Ladau

## Abstract

We examined hypotheses of spatial association arising from direct or indirect competitive interactions, using thirteen years of gridded mammal census data from Kruger National Park, South Africa. As interactions occur at different scales, we explored the data at 1, 5, 10, and 15 km^2^. We proposed four hypotheses structuring the mammal community: H1. direct competition between carnivores and herbivores; H2-4: indirect competition produced by each of three types of herbivore diet specialization: H2. ruminants and non-ruminants, H3. grazers and browsers, and H4. a four-way division of small and large grazers and browsers. We used the software CoOccur to implement a robus test for evidence of our hypotheses against null models of community assemblage. At 5, 10, and 15km^2^ scales, the results supported a competition mechanism in the majority of years for hypotheses H1, H3, and H4, and facilitation in H2. At the finest spatial scale (1km^2^), we saw evidence for a mixture of competitive, neutral and facilitative process. These results suggest strong, large-scale effects of interspecific interactions on distributions of African megafauna, which may not operate at a more local (1km^2^) scale, underscoring the importance of scale and mechanism in the guild structure of communities.

## Introduction

Understanding and describing how species interactions structure communities remains a fundamental goal of ecology (Diamond 1975, Connor and Simberloff 1979, Schoener 1983, Schluter 1984, Stone and Roberts 1990, Manly 1995, Durant 1998, Roxburgh and Matsuki 1999, Linnell and Strand 2000, Gotelli 2000, Gotelli and McCabe 2002, Gotelli and Ellison 2002, Sfenthourakis et al. 2006, Ulrich et al. 2017). Understanding community structure requires both a spatial and temporal perspective (Linnell and Strand 2000), but few datasets are available that are expansive in both of these scales, in part due to declining sizes and numbers of remnant intact wild populations. We present an analysis of long-term, multi-scale, spatially explicit census data of savanna mammals (Table 1) in Kruger National Park, South Africa, testing hypotheses of competition that shape the mammalian assemblage on the landscape. Using census data from a large landscape (approximately 20,000 km^2^) over multiple years gives us the opportunity to respond to the call by Linnell and Strand (2000) for more research into temporal and spatial aspects of species co-existence, when discussing diverse guilds and communities. Using thirteen years of data, we could assess whether patterns were consistent through time, and these data also allowed us to examine patterns at multiple scales, to quantify the potential extent of behavioral and ecological processes structuring communties.

**Table 1:**
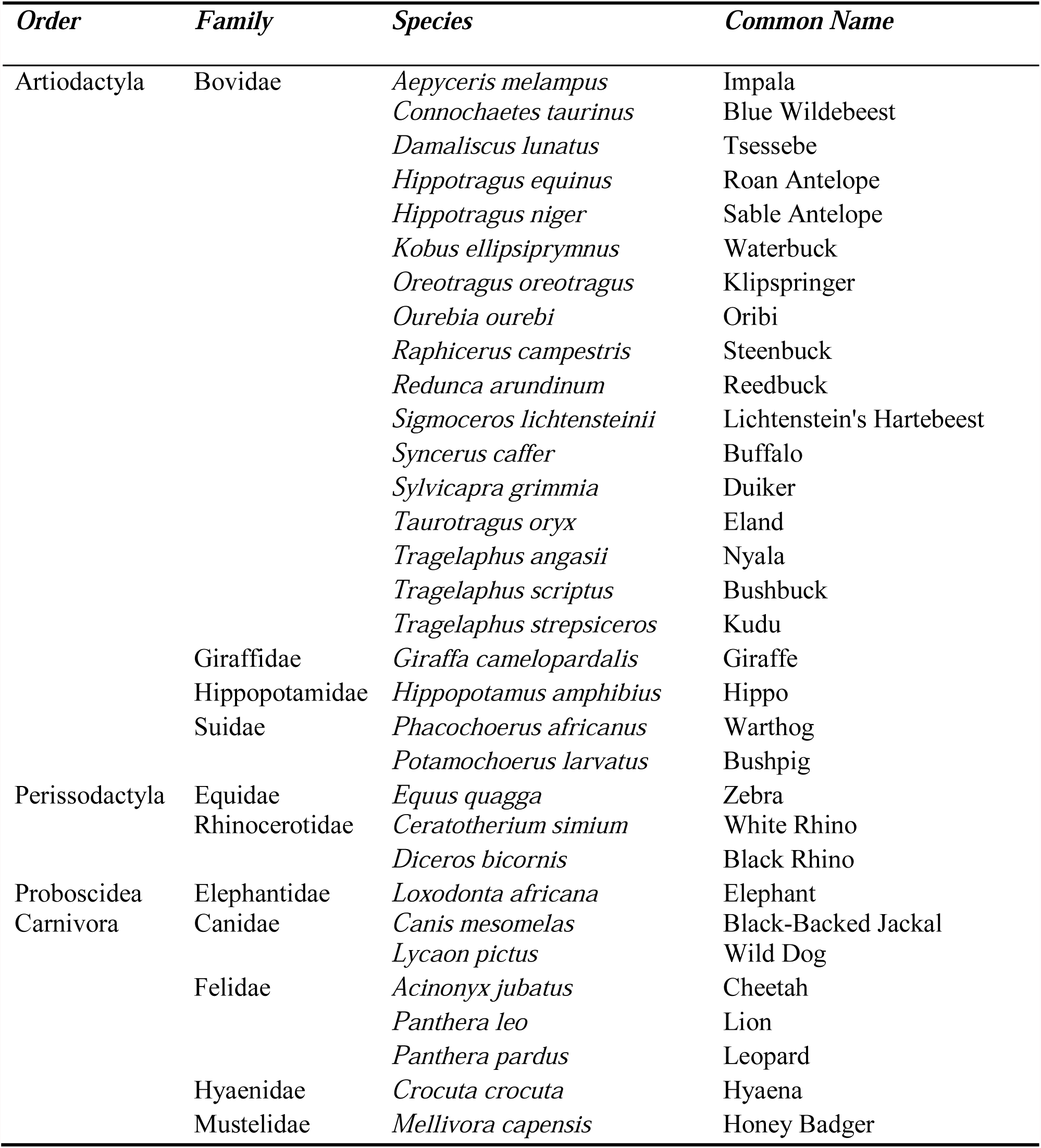
Mammal species observed in the EAS census data and used in the study

### Savanna community structure

Carnivores are likely to shape savanna communities by acting as top predators (Radloff and Toit 2004, Pringle et al. 2007), excluding their prey through direct competition from predation, or via prey avoidance and vigilance behavior (Schmitz 2007). This is expected to generate patterns of competitive exclusion in presence-absence data. We analyzed data collected on larger carnivores in the Felidae (lion, cheetah, leopard) and Canidae (Black-Backed Jackal, Wild dog), plus the hyena and the honey badger. As the predator trophic level, they are more sparsely represented in the data than their prey, but are nevertheless expected to exert detectable effects on the structure of the community.

For large herbivores, the two dominant processes structuring assemblages in African savannas are facilitation and competition (Arsenault and Owen-Smith 2002). Interspecific competition predominates during the dry season, while plants are dormant and high quality forage becomes depleted. Therefore, it has been suggested that in the dry season, this will lead to competition among species with similar requirements, subject to habitat heterogeneity at different scales (Redfern et al. 2006). As these census data were collected during the dry season, we expect a dominance of interspecific competition to be occurring. At large scales, abiotic characteristics of the landscape, such as underlying geology, will affect herbivore distribution (Bailey et al. 1996). At smaller scales, biotic factors such as forage quality and quantity will determine foraging decisions. The selection of feeding ranges, patches, plant species, and even plant parts, will occur at multiple scales (Senft et al. 1987, Senft 1989). Arsenault and Owen-Smith (2002) suggested that facilitation occurs primarily during the plant growing season, as foraging by larger species can create a temporary improvement of forage conditions for smaller species.

However, the spatial co-occurrence of herbivores may also result from habitat selection – species feeding in the same area – rather than causal facilitation. Landscape configuration may thus generate patterns of apparent facilitation, depending on the scale of selection. Distinguishing behavioral differences between these two processes, which would produce identical patterns in the data, requires direct observation and different methods than those we use here.

Based on existing theories of competition and community structure, we developed four hypotheses (H1-4) of mechanisms generating patterns of competition for the vertebrate community of Kruger National Park.

H1. We hypothesize that carnivores exclude their herbivore prey, either through direct predation, or prey avoidance strategies (trait-mediated indirect effects), showing competitive (a lack of co-occurrence) patterns at multiple scales.

H2-4: Within the herbivore suite, we hypothesize that interference competition structures the assemblage via the ability to process food, leading to indirect competition through resource manipulation or interspecific competition for similar diet or habitat (de Boer and Prins 1990).

H2: There is a guild division between ruminants and non-ruminants, based on the premise that gut morphology and digestive ability determine the ability to exploit certain habitats. We thus expect that each guild can exploit different habitats, leading to less co-occurrence of guilds, and a pattern consistent with competition.

H3: A smaller suite of herbivores with a higher degree of habitat overlap represent a guild division between grazers (Buffalo, Zebra, Wildebeest) and browsers (Elephant, Giraffe, Kudu, Impala). Grazers and browsers can putatively exploit different food types, leading to less spatial co-occurrence. While elephants are capable of grazing and browsing, since these data are primarily dry season occurrences, we expect to see elephant occurrence reflecting a browsing strategy. These herbivores will compete within feeding type (guild), although not necessarily between feeding types, leading to indirect competition and generating a pattern consistent with competition.

H4: Competitive structure within the assemblage can arise through a combination of feeding type and body size. This generates indirect competition through highly specialized feeding strategies. Four guild divisions are designated: 1. Large grazers (Buffalo) 2. Small grazers (Zebra and Wildebeest) 3. Large browsers (Elephant) 4. Small browsers (Giraffe, Kudu, Impala) (7 species), following Redfern et al. (2006) (and see Table 2). Feeding type and body size will affect competitive ability for resources, and this will likely occur between different species at different spatial scales, and generate a pattern consistent with competition.

**Table 2:** Species divisions into guilds as specified in hypotheses H1-H4.

| *Hypothesis* | *Guild divisions and species (common names)* |  |
| --- | --- | --- |
| **H1** | **Carnivores:**<br>Black-Backed Jackal, Wild dog,<br>Cheetah, Lion, Leopard, Hyaena,<br>Honey Badger | **Herbivores:**<br>Impala, Blue Wildebeest, Tsessebe, Roan<br>Antelope, Sable Antelope, Waterbuck,<br>Klipspringer, Oribi, Steenbuck, Reedbuck,<br>Lichtenstein's Hartebeest, Buffalo, Duiker,<br>Eland, Nyala, Bushbuck, Kudu, Giraffe,<br>Hippo, Warthog, Bushpig, Zebra, White<br>Rhino, Black Rhino, Elephant |
| **H2** | **Ruminants:**<br>Roan Antelope Bull, Buffalo,<br>Bushbuck, Duiker, Eland, Kudu,<br>Giraffe, Klipspringer,<br>Lichtenstein's Hartebeest, Nyala,<br>Oribi, Impala, Roan Antelope,<br>Reedbuck, Sable Antelope,<br>Steenbuck, Tsessebe, Blue<br>Wildebeest, Waterbuck | **Non-Ruminants:**<br>Bushpig, Hippo, Zebra, Elephant, Black<br>Rhino, Warthog, White Rhino |
| **H3** | **Grazers:**<br>Buffalo, Zebra, Wildebeest | **Browsers:**<br>Elephant, Giraffe, Kudu, Impala |
| **H4** | **Grazers:**<br>*Small*<br>Zebra, Wildebeest<br>*Large*<br>Buffalo | **Browsers:**<br>*Small*<br>Giraffe, Kudu, Impala<br>*Large*<br>Elephant |

### Models of community assemblage

The underlying ecological mechanisms and rules of community assemblage have been the subject of studies in multiple systems (Diamond 1975, Connor and Simberloff 1979, Schoener 1983, Schluter 1984, Stone and Roberts 1990, Manly 1995, Durant 1998, Roxburgh and Matsuki 1999, Linnell and Strand 2000, Gotelli 2000, Gotelli and McCabe 2002, Gotelli and Ellison 2002, Sfenthourakis et al. 2006). Quantifying these patterns, or lack thereof, has led to a long debate in the literature (reviewed by Gotelli and McCabe 2002), sparked by an initial study by Diamond (1975) positing that species assemblages occur through competitive interactions and rebutted by Connor and Simberloff four years later (1979), who demonstrated that random events of colonization could produce similar co-occurrence patterns.

Connor and Simberloff’s (1979) initial introduction of null models of community structure was motivated by an interest to understand which mechanisms (e.g., competition) were responsible for co-occurrence patterns. We follow this approach, developing null model tests based on our specific hypotheses of the ecological mechanism of competition shaping the community structure in question. We use a null model, developed by Ladau and Schwager (2008), that yields a prediction assuming only an absence of competitive interactions (i.e., that species occur independently of each other). Hence, finding data that are inconsistent with this model allows a strong inference about whether competition shapes community assembly.

## Methods

### Study site

The Kruger National Park is located in the lowveld of South Africa, with a core area of nearly 20,000km^2^, at the time of data collection. It has a north-south rainfall gradient of around 440mm-750mm and is dominated by four major landscapes, essentially dividing the park into quadrants due to a basalt/granite geological split the length of the park (Gertenbach 1980, 1983, Redfern et al. 2006) (Figure 1).

**Figure 1:**
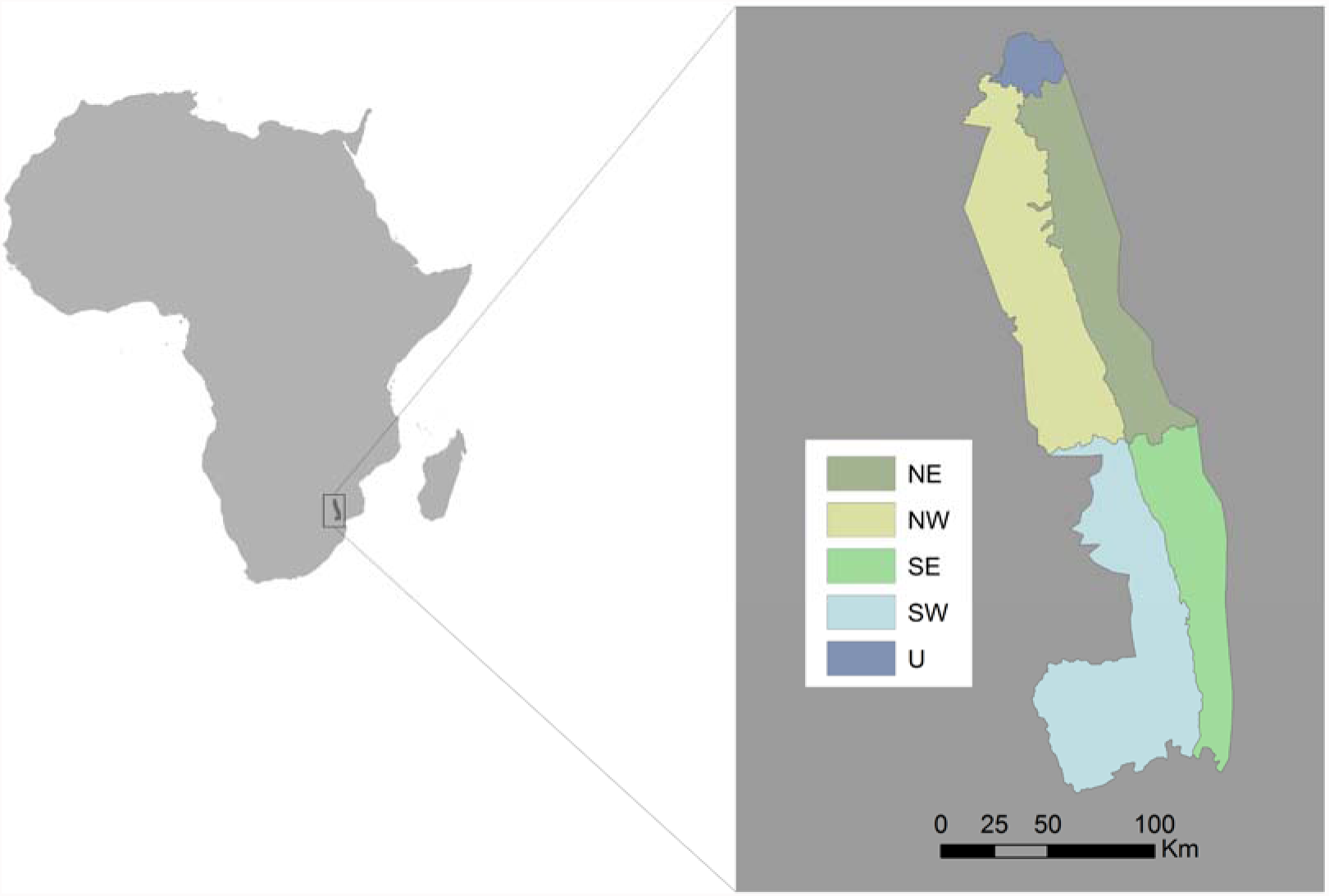
Location of Kruger National Park in Africa, with inset showing the four quadrants (NE, NW, SE, SW) created by the East-West granitic-basaltic landscape split, and the North-South division as a proxy for rain gradient, and a far northern, uncensused (U) region of the park.

### Data

We analyzed the annual fixed-wing census from Kruger National Park collected during 1981-1993. The Ecological Aerial Survey (EAS) was conducted annually, during the dry season between May and August, using a total area count on 800m-wide strip transects (Viljoen 1996a, 1996b). Point counts of herds or groups of animals (listed in Table 1) were conducted by four observers in the plane, and recorded to spatial location by GPS, using four corners and a center point on a quadrat system approximately 2km on a side (Viljoen 1996a). The observation recording patterns changed between 1984 and 1985, in which the number of positions in a sample grid space recorded was increased from four corners to an additional centroid position (Viljoen 1996a, 1996b). This affects the 1km data structure by increasing the number of grid cells 4-fold between the two years. Although the method suffers population level bias from undercounts (Redfern and Getz 2002), it is a systematic bias, allowing us to make appropriate hypothesis comparisons.

### Data sampling method

We created grids of the park boundary at 1km, 5km, 10km, and 15km resolutions, and aggregated the EAS data species counts to grid cells, removing partial boundary grid cells from the analyses. The resulting data sets comprise 17,813, 621, 130 and 43 grid cells, or sites, respectively. GIS data manipulations were conducted in ArcGIS 10.2. These data were then converted to binary species presence-absence data in R.

### Statistical tests of species associations

The EAS data we analyzed for each hypothesis comprise four subsets, sequentially reducing the number of species (Table 2).

The competition model from which the null model is developed is based on two premises:

I. Regardless of how competition acts – evolutionarily or ecologically, and extrinsically or intrinsically – it will reduce the co-occurrence of ecologically similar organisms, which we refer to as guilds.
II. Ecological similarity can be specified using a hierarchical classification system. Species can be classified into “units,” with species in the same unit being more ecologically similar to each other than those in different units. The quality of “units” can be interpreted, for example, as guilds or trophic levels, depending on the hypothesized structure of competition or interactions.

Defining the ⟨*ij*⟩ as the event that the *i*th and *j*th species to arrive at a community belong to the same unit, competitive structuring predicts

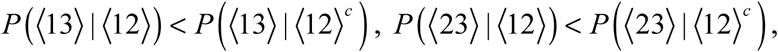

or

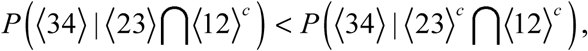

(where c denotes complement). Thus, an appropriate null hypothesis for testing for the absence of interspecific interactions is to set all relevant pairs of conditional probabilities equal. This alone is sufficient to specify a distribution on the set of partitions of species into units, with minimal additional assumptions: there is no autocorrelation, and *P*(⟨*ij*⟩) > 0 for all *i* and *j*.

Hence, to test robustly for interspecific interactions, it is sufficient to assess the agreement between observed and predicted distributions of partitions.

The hypothesis test that we used was as follows:

1. At each site, in this case, a grid cell, the partition of species into units was computed. For example, if three species were observed, two in the same unit (guild) and the other in a different one, then the partition would be “2-1.” If four species were observed, all in different units (guilds), the partition would be “1-1-1-1.”
2. We computed the total sum of squares of the block sizes across all partitions. In the above example, the sum would be 2^2^ + 1^2^ + 1^2^ + 1^2^ + 1^2^ + 1^2^ = 9. We used this sum as our test statistic (*V*).
3. Conditional on the observed number of species (*c*) and units (*u*), the probability of a given partition *I* is *ψ*(*I*) *S*(*c*,*u*), where *ψ*(*I*) gives the number of set partitions corresponding to the given species-unit partition, and *S* is a Stirling number of the second kind - the number of possible ways to partition *n* elements into *k* non-empty subsets. Conditional on the number of species observed at each site, we used this result to compute the expectation value and variance of the sum of squares at each site.
4. With the computed expectation values and variances for each site, we used a 2-tailed test for patterns of both competition and facilitation (α=0.05, Lyapunov CLT). We also used a one-tailed version (Ladau and Schwager 2008), which has power to reject the null hypothesis only if the deviation from it is consistent with effects of competition. This test is uniformly most powerful (UMP) against a broad range of alternatives consistent with competition. We also numerically confirmed that tests had adequate power (>0.80).

We implemented this test using software “CoOccur” written in Visual Basic 6.0, developed by the authors, and available online (http://www.sadieryan.net/cooccur.html). We summarized the results for each hypothesis in Table 3, and report the significance, size and power of the one- and two-tailed tests in full in Supplemental Table 1.

**Table 3:** Summary results of the robust test, by mechanism (H1-H4) and resolution; number of years are given in their *p*-value ranges

| *Mechanism* | *Spatial Resolution* | *p* < 0.05 | 0.05 ≤ *p* ≤ 0.95 | *p* > 0.95 |
| --- | --- | --- | --- | --- |
| H1 | 1km | 2 | 5 | 6 |
|  | 5 km | 13 | -- | -- |
|  | 10 km | 13 | -- | -- |
|  | 15 km | 8 | 5 | -- |
| H2 | 1km | -- | -- | 13 |
|  | 5 km | -- | -- | 13 |
|  | 10 km | -- | -- | 13 |
|  | 15 km | -- | -- | 13 |
| H3 | 1km | 12 | 1 | -- |
|  | 5 km | 13 | -- | -- |
|  | 10 km | 13 | -- | -- |
|  | 15 km | 13 | -- | -- |
| H4 | 1km | 9 | 4 | -- |
|  | 5 km | 8 | 3 | 2 |
|  | 10 km | 13 | -- | -- |
|  | 15 km | 13 | -- | -- |

## Results

The results varied by both scale and hypothesis in question. Results for potential predation effects (H1), at both 5km and 10km, suggested competition in all years (*p*<0.05) and 8 of 13 years at15km resolution (*p*<0.05). At 1km resolution, 2 years suggested competition (*p*<0.05); six years suggested facilitation (*p*=1.00) and the other years were not indicative of a mechanism. At 1km, the one-sided test only had sufficient size and power (minimum and maximum) for the first 4 years; the two-tailed test had sufficient size for these four years and for two additional years, but had insufficient power after the first four years. We thus cannot make robust conclusions after the first four years at the 1km scale.

Perhaps due to sparse observations of carnivores due to their low abundance, or due to the change in recording methods for observations, we found that the minimum criterion of ([n genera + 2]* species) at a site was not always met, drastically reducing the power of the test for H1. This meant that tests at 1km, after 1984, had no power, and therefore did not produce reliable answers, including the 6 years’ results suggesting facilitation. For the remaining 44 tests for H1, all the one-tailed tests had sufficient size and power, as did 43 of the two-tailed tests (Table 3, Table S1).

The ruminant and non-ruminant hypothesis (H2) for structure arising from competition, generated results indicating facilitation *p*=1.00 at all scales in all years. The size and power of all tests for this mechanism was sufficient. For feeding competition between grazers and browsers (H3), all years and scales indicated competition (*p*<0.05), except 1991 at 1km where *p*=0.05. All tests for this mechanism, both one and two-tailed, had sufficient size and power (see Table 3, Table S1).

H4 considered the contributions of grazing and browsing and also body size to competition; the test indicated competition in 9 of the years at a 1km resolution. At a 5km resolution, the test suggested competition in 8 of the years, and facilitation in 2 of the years. At both 10km and 15km resolution, competition was suggested by the test in all 13 years. The size and minimum and maximum power criteria were satisfied for both the one and two-tailed tests in all cases (Table 3, Table S1).

In all cases (4 mechanisms, 13 years, 4 resolutions; total: 208 cases), the Type I error rate of both the one and two-tailed tests were acceptably low. In addition, the power was sufficiently high for all cases, both one and two-tailed tests, with the exception of years 1985-1993 at 1km resolution for H1 (Table 3, Table S1).

## Discussion

We sought to test hypotheses describing the mechanisms that may structure the community of savanna mammals on the Kruger National Park landscape, using tests for species assemblage that allow strong conclusions about mechanisms. We found support for competitive structure patterning for three of our four hypotheses, while the other suggested facilitation in all years.

Testing suites of species involved in specific guild structuring mechanisms meant we could explicitly formulate hypotheses. In addition, creating a software interface (CoOccur), in which the size and power of the test is explicitly evaluated, allowed us to explore potential pitfalls in attaining results, but thereby gave us confidence in our conclusions. For the one-tailed test, rejecting the null hypothesis indicates effects of competition (negative non-independence), while rejecting the null hypothesis in the two-tailed test can indicate competition or facilitation (positive non-independence).

We found evidence of competition at the 5km and 10km scales when carnivores were hypothesized to generate direct competition. We only found evidence for competitive structure for roughly 60% of the years in question at the largest scale (15km). A close examination of the years in question (Table 4) reveals no apparent temporal pattern, suggesting that this may be a sufficiently large spatial scale that direct competition may not always exert a strong effect. At the other extreme, at 1km resolution, for the four years with sufficient power, we found evidence for both competition and a lack of interactions. Evidence of direct competition effects via carnivore presence could indicate direct exclusion by predation, but more likely, behavioral avoidance of carnivores by prey species. Predator vigilance and avoidance has been cited as likely shaping herbivore distributions for savanna mammals, even resulting in unpredictable apparent facilitation of herbivores, wherein mixed grazer herds group together (de Boer and Prins 1990).

In contrast to our expectation that the suite of herbivores might show evidence of resource competition based on the division of ruminants and non-ruminants (H2), we found evidence for facilitation at all scales, including the finer 1km scale. It is plausible that the test was instead detecting habitat selection, at least at large scales, consistent with apparent facilitation due to habitat selection. The ability to monopolize feeding patches at a scale relevant to these herbivores, in terms of home range or daily traveling distance (Estes 1991) (1km or 5km), appears not to be conferred by ruminant ability. However, the more specialized herbivore suites, with guilds of grazers and browsers and even small and large body size (H3 and H4) showed patterns consistent with competitive structure (except one at 1km resolution). The grazer/browser guild showed competition at 5km, 10km, and 15km, and inclusion of body size as a factor also showed patterns consistent with competition at 10km and 15km resolutions, but was less consistent at 5km and 10km. As pointed out by Farnsworth et al. (2002), in theory, organisms sharing the same niche should competitively exclude each other to the point of one species going extinct. However, a model of specialization by plant parts suggests that grazers can co-exist on a landscape, if they can exploit different parts of plant structure. We found that if we partition the guilds along the lines of niche specialization, i.e. grazers and browsers, we start to find evidence of competitive structure. It is plausible that the very broad guilds of ruminants and non-ruminants show apparent facilitation due to clumping of herbivores on the landscape into preferred habitats.

Fritz (1997) conducted a comprehensive study on ungulate biomass at multiple sites in Africa to assess whether the patterns in these communities could be attributable to missing predator guilds, or missing megaherbivores. These guilds both exert pressure on other sympatric guilds, but in slightly different ways, leading to interesting feedbacks on the relative abundance and biomass of medium-sized ungulates. In the study, he attributed the missing guild structure in part to high hunting pressures in certain areas. This is a major consideration for reserve design and conservation of these savanna suites of mammals, and the effects of assemblage disruption and community shifts are evident. Feeley (2003), in a study of avian assemblages on islands resulting from establishment of a large reservoir, pointed out that community structure is an important consideration in reserve design. More recently, studies in savanna systems have shown that the removal of large herbivores can cause unexpected trophic cascades, down to the level of complex feedbacks on ant-plant interactions (Pringle et al. 2007, Palmer et al. 2008, Estes et al. 2011).

Current distribution patterns of herbivores on landscapes may result from currently absent or extinct competitors that once shaped the context for current species. This concept, the “ghost of competition” (Connell 1980), is complicated to address, even with the luxury of this type of long term data. In this study, we assume that patterns reflect interactions within the complement of species over a period of thirteen years. Codron et al. (2008) asked whether large-scale climatic factors or niche specialization may have shaped the assemblage of savanna species over much longer timescales (the Quaternary Land Mammal Ages), and found that adaptation through diet specialization (grazing and browsing) appears to continue through recent evolution, and may be plastic throughout evolutionary history.

These census data do not represent a complete inventory of Kruger National Park. In particular, the carnivore suite suffers not only from detection issues, but also omission of several smaller carnivores. With more detailed carnivore behavioral data, we would be able to explore potential facilitative mechanisms which have been suggested for other systems, such as smaller carnivores (hyenas, jackals, etc.) benefiting from behaving as scavengers at larger carnivore (lion, leopard, e.g.) kills (Grange and Duncan 2006). In addition, there were population declines in several of the herbivore species during this time span, attributed in part to high rates of lion predation (Ogutu and Owen-Smith 2003, Owen-Smith et al. 2005). How this might have affected the data is unclear; we found no changes in potential structuring mechanisms within the herbivores.

## Acknowledgments

This work was supported by a Santa Fe Institute Postdoctoral Fellowship and Gordon and Betty Moore Foundation Grant (J.L.), and in part by an NSF Biological Informatics Fellowship (S.J.R.) and funding from the National Center for Ecological Analysis and Synthesis, a Center funded by NSF (Grant EF-0553768), the University of California, Santa Barbara, and the State of California. The authors thank Scientific Services at Kruger National Park for all their support and data access, and we thank AW and EW for comments and edits.

---

**Supplemental Table 1:** Full results for the statistical tests, by hypothesis H1-H4

**Supp 1.**
| **H1** |  | **One-tailed test** |  |  |  | **Two-tailed test** |  |  |  |  |  |
| --- | --- | --- | --- | --- | --- | --- | --- | --- | --- | --- | --- |
| **Scale** | **Year** | **P-value** | **Size** | **Minimum Power** | **Maximum Power** | **P-value** | **Size** | **Minimum Power** | **Maximum Power** | **Rows** | **Cols** |
| 1km | 1981 | 0.08 | 0.04 | 0.07 | 0.22 | 0.15 | 0.05 | 0.04 | 1.00 | 32 | 3377 |
|  | 1982 | 0.00 | 0.04 | 0.07 | 0.31 | 0.00 | 0.05 | 0.05 | 1.00 | 32 | 3347 |
|  | 1983 | 0.06 | 0.05 | 0.06 | 0.22 | 0.12 | 0.04 | 0.04 | 1.00 | 34 | 3366 |
|  | 1984 | 0.00 | 0.04 | 0.07 | 0.26 | 0.00 | 0.05 | 0.05 | 1.00 | 33 | 3390 |
|  | 1985 | 0.12 | 0.01 | 0.00 | 0.00 | 0.23 | 0.05 | 0.00 | 0.29 | 35 | 12402 |
|  | 1986 | 1.00 | 0.00 | 0.00 | 0.00 | 1.00 | 0.00 | 0.00 | 0.00 | 36 | 12234 |
|  | 1987 | 1.00 | 0.00 | 0.00 | 0.00 | 1.00 | 0.00 | 0.00 | 0.00 | 36 | 12493 |
|  | 1988 | 0.16 | 0.00 | 0.00 | 0.00 | 0.32 | 0.05 | 0.00 | 0.30 | 35 | 12351 |
|  | 1989 | 1.00 | 0.00 | 0.00 | 0.00 | 1.00 | 0.00 | 0.00 | 0.00 | 35 | 11718 |
|  | 1990 | 1.00 | 0.00 | 0.00 | 0.00 | 1.00 | 0.00 | 0.00 | 0.00 | 34 | 11905 |
|  | 1991 | 0.12 | 0.00 | 0.00 | 0.00 | 0.25 | 0.00 | 0.00 | 0.00 | 36 | 11330 |
|  | 1992 | 1.00 | 0.00 | 0.00 | 0.00 | 1.00 | 0.00 | 0.00 | 0.00 | 36 | 11811 |
|  | 1993 | 1.00 | 0.00 | 0.00 | 0.00 | 1.00 | 0.00 | 0.00 | 0.00 | 36 | 10772 |
| 5km | 1981 | 0.00 | 0.04 | 0.12 | 0.61 | 0.00 | 0.05 | 0.05 | 1.00 | 31 | 604 |
|  | 1982 | 0.00 | 0.04 | 0.14 | 0.78 | 0.00 | 0.05 | 0.05 | 1.00 | 32 | 605 |
|  | 1983 | 0.00 | 0.04 | 0.13 | 0.77 | 0.00 | 0.05 | 0.05 | 1.00 | 34 | 605 |
|  | 1984 | 0.00 | 0.05 | 0.16 | 0.83 | 0.00 | 0.05 | 0.05 | 1.00 | 32 | 604 |
|  | 1985 | 0.00 | 0.04 | 0.16 | 0.85 | 0.00 | 0.05 | 0.05 | 1.00 | 34 | 617 |
|  | 1986 | 0.00 | 0.04 | 0.13 | 0.73 | 0.00 | 0.05 | 0.05 | 1.00 | 36 | 607 |
|  | 1987 | 0.00 | 0.04 | 0.13 | 0.75 | 0.00 | 0.06 | 0.06 | 1.00 | 35 | 617 |
|  | 1988 | 0.00 | 0.05 | 0.12 | 0.66 | 0.00 | 0.05 | 0.05 | 1.00 | 35 | 619 |
|  | 1989 | 0.00 | 0.04 | 0.11 | 0.53 | 0.00 | 0.05 | 0.05 | 1.00 | 35 | 620 |
|  | 1990 | 0.00 | 0.04 | 0.13 | 0.71 | 0.00 | 0.05 | 0.05 | 1.00 | 34 | 611 |
|  | 1991 | 0.01 | 0.04 | 0.09 | 0.46 | 0.02 | 0.05 | 0.05 | 1.00 | 36 | 620 |
|  | 1992 | 0.00 | 0.04 | 0.09 | 0.37 | 0.00 | 0.05 | 0.05 | 1.00 | 35 | 621 |
|  | 1993 | 0.01 | 0.03 | 0.06 | 0.18 | 0.02 | 0.05 | 0.01 | 1.00 | 36 | 620 |

| H1 |  | One-tailed test |  |  |  | Two-tailed test |  |  |  |  |  |
| --- | --- | --- | --- | --- | --- | --- | --- | --- | --- | --- | --- |
| Scale | Year | P-value | Size | Minimum Power | Maximum Power | P-value | Size | Minimum Power | Maximum Power | Rows | Cols |
| 10km | 1981 | 0.00 | 0.04 | 0.13 | 0.68 | 0.00 | 0.04 | 0.04 | 1.00 | 31 | 130 |
|  | 1982 | 0.00 | 0.04 | 0.15 | 0.81 | 0.00 | 0.05 | 0.05 | 1.00 | 32 | 129 |
|  | 1983 | 0.00 | 0.04 | 0.15 | 0.87 | 0.00 | 0.05 | 0.05 | 1.00 | 33 | 130 |
|  | 1984 | 0.00 | 0.05 | 0.17 | 0.88 | 0.00 | 0.05 | 0.05 | 1.00 | 32 | 130 |
|  | 1985 | 0.00 | 0.05 | 0.16 | 0.88 | 0.01 | 0.05 | 0.05 | 1.00 | 34 | 130 |
|  | 1986 | 0.00 | 0.04 | 0.14 | 0.81 | 0.00 | 0.05 | 0.05 | 1.00 | 36 | 130 |
|  | 1987 | 0.00 | 0.04 | 0.15 | 0.81 | 0.00 | 0.05 | 0.05 | 1.00 | 35 | 130 |
|  | 1988 | 0.00 | 0.05 | 0.14 | 0.74 | 0.00 | 0.05 | 0.05 | 1.00 | 35 | 130 |
|  | 1989 | 0.00 | 0.04 | 0.13 | 0.66 | 0.00 | 0.06 | 0.06 | 1.00 | 34 | 130 |
|  | 1990 | 0.00 | 0.04 | 0.14 | 0.75 | 0.00 | 0.05 | 0.05 | 1.00 | 34 | 130 |
|  | 1991 | 0.02 | 0.04 | 0.12 | 0.58 | 0.03 | 0.05 | 0.05 | 1.00 | 36 | 130 |
|  | 1992 | 0.00 | 0.05 | 0.12 | 0.60 | 0.00 | 0.05 | 0.05 | 1.00 | 35 | 130 |
|  | 1993 | 0.00 | 0.03 | 0.08 | 0.40 | 0.00 | 0.04 | 0.04 | 1.00 | 36 | 130 |
| 15km | 1981 | 0.03 | 0.04 | 0.10 | 0.45 | 0.07 | 0.05 | 0.05 | 1.00 | 31 | 43 |
|  | 1982 | 0.00 | 0.04 | 0.11 | 0.56 | 0.00 | 0.05 | 0.05 | 1.00 | 32 | 43 |
|  | 1983 | 0.07 | 0.04 | 0.13 | 0.71 | 0.13 | 0.06 | 0.06 | 1.00 | 33 | 43 |
|  | 1984 | 0.01 | 0.04 | 0.13 | 0.63 | 0.01 | 0.05 | 0.05 | 1.00 | 31 | 43 |
|  | 1985 | 0.11 | 0.04 | 0.12 | 0.62 | 0.21 | 0.05 | 0.05 | 1.00 | 34 | 43 |
|  | 1986 | 0.14 | 0.04 | 0.12 | 0.61 | 0.28 | 0.04 | 0.04 | 1.00 | 36 | 43 |
|  | 1987 | 0.05 | 0.04 | 0.12 | 0.57 | 0.10 | 0.04 | 0.04 | 1.00 | 35 | 43 |
|  | 1988 | 0.01 | 0.03 | 0.11 | 0.53 | 0.01 | 0.05 | 0.05 | 1.00 | 34 | 43 |
|  | 1989 | 0.00 | 0.04 | 0.10 | 0.47 | 0.01 | 0.05 | 0.05 | 1.00 | 34 | 43 |
|  | 1990 | 0.00 | 0.04 | 0.11 | 0.48 | 0.00 | 0.05 | 0.05 | 1.00 | 34 | 43 |
|  | 1991 | 0.12 | 0.04 | 0.09 | 0.43 | 0.24 | 0.04 | 0.04 | 1.00 | 36 | 43 |
|  | 1992 | 0.00 | 0.04 | 0.10 | 0.45 | 0.00 | 0.05 | 0.05 | 1.00 | 35 | 43 |
|  | 1993 | 0.00 | 0.04 | 0.08 | 0.29 | 0.00 | 0.05 | 0.05 | 1.00 | 35 | 43 |

| H2 |  | One-tailed test |  |  |  | Two-tailed test |  |  |  |  |  |
| --- | --- | --- | --- | --- | --- | --- | --- | --- | --- | --- | --- |
| Scale | Year | P-value | Size | Minimum Power | Maximum Power | P-value | Size | Minimum Power | Maximum Power | Rows | Cols |
| 1km | 1981 | 1.00 | 0.05 | 0.00 | 1.00 | 0.00 | 0.05 | 0.04 | 1.00 | 26 | 3377 |
|  | 1982 | 1.00 | 0.05 | 0.00 | 1.00 | 0.00 | 0.05 | 0.04 | 1.00 | 27 | 3347 |
|  | 1983 | 1.00 | 0.05 | 0.01 | 1.00 | 0.00 | 0.05 | 0.05 | 1.00 | 28 | 3366 |
|  | 1984 | 1.00 | 0.06 | 0.00 | 1.00 | 0.00 | 0.06 | 0.03 | 1.00 | 27 | 3390 |
|  | 1985 | 1.00 | 0.06 | 0.08 | 1.00 | 0.00 | 0.06 | 0.06 | 1.00 | 29 | 12402 |
|  | 1986 | 1.00 | 0.05 | 0.07 | 1.00 | 0.00 | 0.06 | 0.06 | 1.00 | 29 | 12234 |
|  | 1987 | 1.00 | 0.07 | 0.08 | 1.00 | 0.00 | 0.05 | 0.05 | 1.00 | 29 | 12493 |
|  | 1988 | 1.00 | 0.05 | 0.07 | 1.00 | 0.00 | 0.05 | 0.05 | 1.00 | 28 | 12351 |
|  | 1989 | 1.00 | 0.05 | 0.06 | 1.00 | 0.00 | 0.05 | 0.05 | 1.00 | 28 | 11718 |
|  | 1990 | 1.00 | 0.05 | 0.06 | 1.00 | 0.00 | 0.05 | 0.05 | 1.00 | 28 | 11905 |
|  | 1991 | 1.00 | 0.06 | 0.07 | 1.00 | 0.00 | 0.06 | 0.06 | 1.00 | 29 | 11330 |
|  | 1992 | 1.00 | 0.05 | 0.07 | 0.99 | 0.00 | 0.05 | 0.05 | 1.00 | 29 | 11811 |
|  | 1993 | 1.00 | 0.05 | 0.07 | 0.97 | 0.00 | 0.05 | 0.05 | 1.00 | 30 | 10772 |
| 5km | 1981 | 1.00 | 0.05 | 0.00 | 1.00 | 0.00 | 0.06 | 0.06 | 1.00 | 26 | 604 |
|  | 1982 | 1.00 | 0.04 | 0.00 | 1.00 | 0.00 | 0.04 | 0.04 | 1.00 | 27 | 605 |
|  | 1983 | 1.00 | 0.05 | 0.00 | 1.00 | 0.00 | 0.06 | 0.06 | 1.00 | 28 | 605 |
|  | 1984 | 1.00 | 0.05 | 0.00 | 1.00 | 0.00 | 0.05 | 0.05 | 1.00 | 27 | 604 |
|  | 1985 | 1.00 | 0.04 | 0.00 | 1.00 | 0.00 | 0.05 | 0.05 | 1.00 | 29 | 617 |
|  | 1986 | 1.00 | 0.05 | 0.00 | 1.00 | 0.00 | 0.05 | 0.05 | 1.00 | 29 | 607 |
|  | 1987 | 1.00 | 0.05 | 0.00 | 1.00 | 0.00 | 0.05 | 0.05 | 1.00 | 29 | 617 |
|  | 1988 | 1.00 | 0.05 | 0.00 | 1.00 | 0.00 | 0.05 | 0.05 | 1.00 | 28 | 619 |
|  | 1989 | 1.00 | 0.06 | 0.00 | 1.00 | 0.00 | 0.06 | 0.06 | 1.00 | 28 | 620 |
|  | 1990 | 1.00 | 0.06 | 0.00 | 1.00 | 0.00 | 0.06 | 0.06 | 1.00 | 28 | 611 |
|  | 1991 | 1.00 | 0.05 | 0.00 | 1.00 | 0.00 | 0.05 | 0.05 | 1.00 | 29 | 620 |
|  | 1992 | 1.00 | 0.05 | 0.00 | 1.00 | 0.00 | 0.05 | 0.05 | 1.00 | 29 | 621 |
|  | 1993 | 1.00 | 0.05 | 0.00 | 1.00 | 0.00 | 0.05 | 0.05 | 1.00 | 30 | 620 |
| Scale | Year | P-value | Size | Minimum Power | Maximum Power | P-value | Size | Minimum Power | Maximum Power | Rows | Cols |
|  | 1982 | 1.00 | 0.04 | 0.00 | 1.00 | 0.00 | 0.05 | 0.04 | 1.00 | 27 | 129 |
|  | 1983 | 1.00 | 0.04 | 0.00 | 1.00 | 0.00 | 0.05 | 0.02 | 1.00 | 28 | 130 |
|  | 1984 | 1.00 | 0.04 | 0.00 | 1.00 | 0.00 | 0.05 | 0.05 | 1.00 | 27 | 130 |
|  | 1985 | 1.00 | 0.04 | 0.00 | 1.00 | 0.00 | 0.05 | 0.04 | 1.00 | 29 | 130 |
|  | 1986 | 1.00 | 0.05 | 0.00 | 1.00 | 0.00 | 0.05 | 0.03 | 1.00 | 29 | 130 |
|  | 1987 | 1.00 | 0.05 | 0.00 | 1.00 | 0.00 | 0.05 | 0.02 | 1.00 | 29 | 130 |
|  | 1988 | 1.00 | 0.05 | 0.00 | 1.00 | 0.00 | 0.06 | 0.02 | 1.00 | 28 | 130 |
|  | 1989 | 1.00 | 0.04 | 0.00 | 1.00 | 0.00 | 0.04 | 0.02 | 1.00 | 28 | 130 |
|  | 1990 | 1.00 | 0.04 | 0.00 | 1.00 | 0.00 | 0.05 | 0.03 | 1.00 | 28 | 130 |
|  | 1991 | 1.00 | 0.05 | 0.00 | 1.00 | 0.00 | 0.05 | 0.01 | 1.00 | 29 | 130 |
|  | 1992 | 1.00 | 0.05 | 0.00 | 1.00 | 0.00 | 0.05 | 0.01 | 1.00 | 29 | 130 |
|  | 1993 | 1.00 | 0.05 | 0.00 | 1.00 | 0.00 | 0.06 | 0.02 | 1.00 | 30 | 130 |
| 15km | 1981 | 1.00 | 0.04 | 0.00 | 1.00 | 0.00 | 0.05 | 0.02 | 1.00 | 26 | 43 |
|  | 1982 | 1.00 | 0.03 | 0.00 | 1.00 | 0.00 | 0.05 | 0.02 | 1.00 | 27 | 43 |
|  | 1983 | 1.00 | 0.03 | 0.00 | 1.00 | 0.00 | 0.05 | 0.03 | 1.00 | 28 | 43 |
|  | 1984 | 1.00 | 0.03 | 0.00 | 1.00 | 0.00 | 0.04 | 0.02 | 1.00 | 27 | 43 |
|  | 1985 | 1.00 | 0.04 | 0.00 | 1.00 | 0.00 | 0.05 | 0.03 | 1.00 | 29 | 43 |
|  | 1986 | 1.00 | 0.04 | 0.00 | 1.00 | 0.00 | 0.05 | 0.02 | 1.00 | 29 | 43 |
|  | 1987 | 1.00 | 0.04 | 0.00 | 1.00 | 0.00 | 0.05 | 0.02 | 1.00 | 29 | 43 |
|  | 1988 | 1.00 | 0.04 | 0.00 | 1.00 | 0.00 | 0.05 | 0.02 | 1.00 | 28 | 43 |
|  | 1989 | 1.00 | 0.04 | 0.00 | 1.00 | 0.00 | 0.04 | 0.02 | 1.00 | 28 | 43 |
|  | 1990 | 1.00 | 0.04 | 0.00 | 1.00 | 0.00 | 0.05 | 0.02 | 1.00 | 28 | 43 |
|  | 1991 | 1.00 | 0.04 | 0.00 | 1.00 | 0.00 | 0.05 | 0.02 | 1.00 | 29 | 43 |
|  | 1992 | 1.00 | 0.04 | 0.00 | 1.00 | 0.00 | 0.05 | 0.02 | 1.00 | 29 | 43 |
|  | 1993 | 1.00 | 0.04 | 0.00 | 1.00 | 0.00 | 0.04 | 0.02 | 1.00 | 29 | 43 |

| **H3** |  | **One-tailed test** |  |  |  | **Two-tailed test** |  |  |  |  |  |
| --- | --- | --- | --- | --- | --- | --- | --- | --- | --- | --- | --- |
| **Scale** | **Year** | **P-value** | **Size** | **Minimum Power** | **Maximum Power** | **P-value** | **Size** | **Minimum Power** | **Maximum Power** | **Rows** | **Cols** |
| 1km | 1981 | 0.00 | 0.05 | 0.04 | 1.00 | 0.00 | 0.05 | 0.05 | 1.00 | 11 | 3377 |
|  | 1982 | 0.00 | 0.05 | 0.04 | 1.00 | 0.00 | 0.05 | 0.05 | 1.00 | 11 | 3347 |
|  | 1983 | 0.00 | 0.04 | 0.05 | 1.00 | 0.00 | 0.04 | 0.04 | 1.00 | 11 | 3366 |
|  | 1984 | 0.00 | 0.05 | 0.04 | 1.00 | 0.00 | 0.05 | 0.05 | 1.00 | 11 | 3390 |
|  | 1985 | 0.00 | 0.05 | 0.07 | 0.98 | 0.01 | 0.05 | 0.05 | 1.00 | 11 | 12402 |
|  | 1986 | 0.00 | 0.05 | 0.07 | 0.89 | 0.00 | 0.05 | 0.05 | 0.98 | 11 | 12234 |
|  | 1987 | 0.00 | 0.05 | 0.07 | 0.90 | 0.00 | 0.05 | 0.05 | 0.99 | 11 | 12493 |
|  | 1988 | 0.00 | 0.06 | 0.07 | 0.96 | 0.00 | 0.05 | 0.05 | 1.00 | 11 | 12351 |
|  | 1989 | 0.01 | 0.05 | 0.06 | 0.87 | 0.02 | 0.05 | 0.04 | 0.97 | 11 | 11718 |
|  | 1990 | 0.01 | 0.05 | 0.07 | 0.91 | 0.02 | 0.05 | 0.05 | 0.99 | 11 | 11905 |
|  | 1991 | 0.05 | 0.05 | 0.07 | 0.86 | 0.10 | 0.06 | 0.05 | 0.97 | 11 | 11330 |
|  | 1992 | 0.00 | 0.05 | 0.07 | 0.83 | 0.00 | 0.05 | 0.05 | 0.96 | 11 | 11811 |
|  | 1993 | 0.00 | 0.05 | 0.07 | 0.75 | 0.00 | 0.05 | 0.05 | 0.91 | 11 | 10772 |
| 5km | 1981 | 0.00 | 0.05 | 0.00 | 1.00 | 0.00 | 0.05 | 0.05 | 1.00 | 11 | 604 |
|  | 1982 | 0.00 | 0.05 | 0.00 | 1.00 | 0.00 | 0.05 | 0.05 | 1.00 | 11 | 605 |
|  | 1983 | 0.00 | 0.05 | 0.00 | 1.00 | 0.00 | 0.05 | 0.05 | 1.00 | 11 | 605 |
|  | 1984 | 0.00 | 0.05 | 0.00 | 1.00 | 0.00 | 0.05 | 0.05 | 1.00 | 11 | 604 |
|  | 1985 | 0.00 | 0.05 | 0.00 | 1.00 | 0.00 | 0.05 | 0.05 | 1.00 | 11 | 617 |
|  | 1986 | 0.00 | 0.05 | 0.00 | 1.00 | 0.00 | 0.05 | 0.05 | 1.00 | 11 | 607 |
|  | 1987 | 0.00 | 0.04 | 0.00 | 1.00 | 0.00 | 0.05 | 0.05 | 1.00 | 11 | 617 |
|  | 1988 | 0.00 | 0.05 | 0.00 | 1.00 | 0.00 | 0.05 | 0.05 | 1.00 | 11 | 619 |
|  | 1989 | 0.00 | 0.05 | 0.00 | 1.00 | 0.00 | 0.05 | 0.05 | 1.00 | 11 | 620 |
|  | 1990 | 0.00 | 0.04 | 0.00 | 1.00 | 0.00 | 0.05 | 0.05 | 1.00 | 11 | 611 |
|  | 1991 | 0.00 | 0.05 | 0.00 | 1.00 | 0.00 | 0.05 | 0.05 | 1.00 | 11 | 620 |
|  | 1992 | 0.00 | 0.05 | 0.00 | 1.00 | 0.00 | 0.05 | 0.05 | 1.00 | 11 | 621 |
|  | 1993 | 0.00 | 0.05 | 0.01 | 1.00 | 0.00 | 0.05 | 0.05 | 1.00 | 11 | 620 |
| **Scale** | **Year** | **P-value** | **Size** | **Minimum Power** | **Maximum Power** | **P-value** | **Size** | **Minimum Power** | **Maximum Power** | **Rows** | **Cols** |
|  | 1982 | 0.00 | 0.04 | 0.00 | 1.00 | 0.00 | 0.05 | 0.05 | 1.00 | 11 | 129 |
|  | 1983 | 0.00 | 0.04 | 0.00 | 1.00 | 0.00 | 0.05 | 0.05 | 1.00 | 11 | 130 |
|  | 1984 | 0.00 | 0.05 | 0.00 | 1.00 | 0.00 | 0.05 | 0.05 | 1.00 | 11 | 130 |
|  | 1985 | 0.00 | 0.04 | 0.00 | 1.00 | 0.00 | 0.04 | 0.04 | 1.00 | 11 | 130 |
|  | 1986 | 0.00 | 0.05 | 0.00 | 1.00 | 0.00 | 0.05 | 0.05 | 1.00 | 11 | 130 |
|  | 1987 | 0.00 | 0.05 | 0.00 | 1.00 | 0.00 | 0.05 | 0.05 | 1.00 | 11 | 130 |
|  | 1988 | 0.00 | 0.05 | 0.00 | 1.00 | 0.00 | 0.05 | 0.05 | 1.00 | 11 | 130 |
|  | 1989 | 0.00 | 0.05 | 0.00 | 1.00 | 0.00 | 0.05 | 0.05 | 1.00 | 11 | 130 |
|  | 1990 | 0.00 | 0.04 | 0.00 | 1.00 | 0.00 | 0.05 | 0.05 | 1.00 | 11 | 130 |
|  | 1991 | 0.00 | 0.05 | 0.00 | 1.00 | 0.00 | 0.05 | 0.05 | 1.00 | 11 | 130 |
|  | 1992 | 0.00 | 0.05 | 0.00 | 1.00 | 0.00 | 0.05 | 0.05 | 1.00 | 11 | 130 |
|  | 1993 | 0.00 | 0.04 | 0.00 | 1.00 | 0.00 | 0.05 | 0.05 | 1.00 | 11 | 130 |
| 15km | 1981 | 0.00 | 0.04 | 0.00 | 1.00 | 0.00 | 0.05 | 0.05 | 1.00 | 11 | 43 |
|  | 1982 | 0.00 | 0.04 | 0.00 | 1.00 | 0.00 | 0.04 | 0.04 | 1.00 | 11 | 43 |
|  | 1983 | 0.00 | 0.04 | 0.00 | 1.00 | 0.00 | 0.05 | 0.05 | 1.00 | 11 | 43 |
|  | 1984 | 0.00 | 0.03 | 0.00 | 1.00 | 0.00 | 0.05 | 0.05 | 1.00 | 11 | 43 |
|  | 1985 | 0.00 | 0.04 | 0.00 | 1.00 | 0.00 | 0.05 | 0.05 | 1.00 | 11 | 43 |
|  | 1986 | 0.00 | 0.04 | 0.00 | 1.00 | 0.00 | 0.05 | 0.05 | 1.00 | 11 | 43 |
|  | 1987 | 0.00 | 0.03 | 0.00 | 1.00 | 0.00 | 0.05 | 0.05 | 1.00 | 11 | 43 |
|  | 1988 | 0.00 | 0.03 | 0.00 | 1.00 | 0.00 | 0.05 | 0.05 | 1.00 | 11 | 43 |
|  | 1989 | 0.00 | 0.04 | 0.00 | 1.00 | 0.00 | 0.05 | 0.05 | 1.00 | 11 | 43 |
|  | 1990 | 0.00 | 0.04 | 0.00 | 1.00 | 0.00 | 0.04 | 0.04 | 1.00 | 11 | 43 |
|  | 1991 | 0.00 | 0.04 | 0.00 | 1.00 | 0.00 | 0.05 | 0.05 | 1.00 | 11 | 43 |
|  | 1992 | 0.00 | 0.04 | 0.00 | 1.00 | 0.00 | 0.05 | 0.05 | 1.00 | 11 | 43 |
|  | 1993 | 0.00 | 0.04 | 0.00 | 1.00 | 0.00 | 0.05 | 0.05 | 1.00 | 11 | 43 |

| **H4** |  | **One-tailed test** |  |  |  | **Two-tailed test** |  |  |  |  |  |
| --- | --- | --- | --- | --- | --- | --- | --- | --- | --- | --- | --- |
| **Scale** | **Year** | **P-value** | **Size** | **Minimum Power** | **Maximum Power** | **P-value** | **Size** | **Minimum Power** | **Maximum Power** | **Rows** | **Cols** |
| 1km | 1981 | 0.86 | 0.06 | 0.13 | 1.00 | 0.29 | 0.05 | 0.05 | 1.00 | 11 | 3377 |
|  | 1982 | 0.29 | 0.05 | 0.12 | 1.00 | 0.58 | 0.04 | 0.04 | 1.00 | 11 | 3347 |
|  | 1983 | 0.07 | 0.05 | 0.11 | 1.00 | 0.14 | 0.05 | 0.05 | 1.00 | 11 | 3366 |
|  | 1984 | 0.00 | 0.05 | 0.12 | 1.00 | 0.01 | 0.05 | 0.05 | 1.00 | 11 | 3390 |
|  | 1985 | 0.04 | 0.05 | 0.08 | 0.89 | 0.07 | 0.05 | 0.05 | 0.99 | 11 | 12402 |
|  | 1986 | 0.00 | 0.05 | 0.07 | 0.79 | 0.00 | 0.05 | 0.05 | 0.94 | 11 | 12234 |
|  | 1987 | 0.00 | 0.06 | 0.07 | 0.76 | 0.01 | 0.05 | 0.05 | 0.91 | 11 | 12493 |
|  | 1988 | 0.01 | 0.05 | 0.06 | 0.91 | 0.02 | 0.05 | 0.05 | 0.98 | 11 | 12351 |
|  | 1989 | 0.01 | 0.05 | 0.08 | 0.79 | 0.02 | 0.05 | 0.05 | 0.94 | 11 | 11718 |
|  | 1990 | 0.01 | 0.05 | 0.08 | 0.80 | 0.03 | 0.05 | 0.05 | 0.96 | 11 | 11905 |
|  | 1991 | 0.13 | 0.05 | 0.06 | 0.74 | 0.25 | 0.05 | 0.05 | 0.91 | 11 | 11330 |
|  | 1992 | 0.01 | 0.05 | 0.06 | 0.69 | 0.02 | 0.04 | 0.04 | 0.86 | 11 | 11811 |
|  | 1993 | 0.00 | 0.05 | 0.07 | 0.61 | 0.00 | 0.05 | 0.05 | 0.76 | 11 | 10772 |
| 5km | 1981 | 0.98 | 0.05 | 0.17 | 1.00 | 0.04 | 0.04 | 0.04 | 1.00 | 11 | 604 |
|  | 1982 | 1.00 | 0.05 | 0.21 | 1.00 | 0.00 | 0.04 | 0.04 | 1.00 | 11 | 605 |
|  | 1983 | 0.13 | 0.05 | 0.18 | 1.00 | 0.26 | 0.05 | 0.05 | 1.00 | 11 | 605 |
|  | 1984 | 0.04 | 0.05 | 0.20 | 1.00 | 0.07 | 0.04 | 0.04 | 1.00 | 11 | 604 |
|  | 1985 | 0.42 | 0.05 | 0.21 | 1.00 | 0.84 | 0.05 | 0.05 | 1.00 | 11 | 617 |
|  | 1986 | 0.07 | 0.05 | 0.19 | 1.00 | 0.13 | 0.05 | 0.05 | 1.00 | 11 | 607 |
|  | 1987 | 0.02 | 0.04 | 0.17 | 1.00 | 0.03 | 0.05 | 0.05 | 1.00 | 11 | 617 |
|  | 1988 | 0.00 | 0.05 | 0.20 | 1.00 | 0.00 | 0.05 | 0.05 | 1.00 | 11 | 619 |
|  | 1989 | 0.03 | 0.04 | 0.19 | 1.00 | 0.06 | 0.05 | 0.05 | 1.00 | 11 | 620 |
|  | 1990 | 0.00 | 0.06 | 0.17 | 1.00 | 0.00 | 0.05 | 0.05 | 1.00 | 11 | 611 |
|  | 1991 | 0.00 | 0.04 | 0.16 | 1.00 | 0.00 | 0.05 | 0.05 | 1.00 | 11 | 620 |
|  | 1992 | 0.00 | 0.04 | 0.17 | 1.00 | 0.01 | 0.04 | 0.04 | 1.00 | 11 | 621 |
|  | 1993 | 0.00 | 0.04 | 0.15 | 1.00 | 0.00 | 0.05 | 0.05 | 1.00 | 11 | 620 |
| **Scale** | **Year** | **P-value** | **Size** | **Minimum Power** | **Maximum Power** | **P-value** | **Size** | **Minimum Power** | **Maximum Power** | **Rows** | **Cols** |
|  | 1982 | 0.02 | 0.05 | 0.17 | 1.00 | 0.03 | 0.06 | 0.06 | 1.00 | 11 | 129 |
|  | 1983 | 0.00 | 0.04 | 0.16 | 1.00 | 0.00 | 0.05 | 0.05 | 1.00 | 11 | 130 |
|  | 1984 | 0.00 | 0.04 | 0.17 | 1.00 | 0.00 | 0.05 | 0.05 | 1.00 | 11 | 130 |
|  | 1985 | 0.01 | 0.05 | 0.15 | 1.00 | 0.02 | 0.06 | 0.06 | 1.00 | 11 | 130 |
|  | 1986 | 0.00 | 0.05 | 0.15 | 1.00 | 0.00 | 0.05 | 0.05 | 1.00 | 11 | 130 |
|  | 1987 | 0.00 | 0.05 | 0.16 | 1.00 | 0.00 | 0.05 | 0.05 | 1.00 | 11 | 130 |
|  | 1988 | 0.00 | 0.04 | 0.15 | 1.00 | 0.00 | 0.05 | 0.05 | 1.00 | 11 | 130 |
|  | 1989 | 0.00 | 0.05 | 0.15 | 1.00 | 0.00 | 0.06 | 0.06 | 1.00 | 11 | 130 |
|  | 1990 | 0.00 | 0.05 | 0.16 | 1.00 | 0.00 | 0.05 | 0.05 | 1.00 | 11 | 130 |
|  | 1991 | 0.00 | 0.04 | 0.14 | 1.00 | 0.00 | 0.05 | 0.05 | 1.00 | 11 | 130 |
|  | 1992 | 0.00 | 0.05 | 0.15 | 1.00 | 0.00 | 0.06 | 0.06 | 1.00 | 11 | 130 |
|  | 1993 | 0.00 | 0.05 | 0.13 | 1.00 | 0.00 | 0.05 | 0.05 | 1.00 | 11 | 130 |
| 15km | 1981 | 0.00 | 0.04 | 0.09 | 0.99 | 0.00 | 0.05 | 0.05 | 1.00 | 11 | 43 |
|  | 1982 | 0.00 | 0.04 | 0.09 | 1.00 | 0.00 | 0.05 | 0.05 | 1.00 | 11 | 43 |
|  | 1983 | 0.00 | 0.03 | 0.10 | 1.00 | 0.00 | 0.05 | 0.05 | 1.00 | 11 | 43 |
|  | 1984 | 0.00 | 0.04 | 0.09 | 0.99 | 0.00 | 0.04 | 0.04 | 1.00 | 11 | 43 |
|  | 1985 | 0.00 | 0.04 | 0.09 | 0.99 | 0.00 | 0.04 | 0.04 | 1.00 | 11 | 43 |
|  | 1986 | 0.00 | 0.04 | 0.10 | 0.99 | 0.00 | 0.05 | 0.05 | 1.00 | 11 | 43 |
|  | 1987 | 0.00 | 0.05 | 0.09 | 0.99 | 0.00 | 0.05 | 0.05 | 1.00 | 11 | 43 |
|  | 1988 | 0.00 | 0.05 | 0.10 | 0.99 | 0.00 | 0.05 | 0.05 | 1.00 | 11 | 43 |
|  | 1989 | 0.00 | 0.05 | 0.09 | 0.99 | 0.00 | 0.05 | 0.05 | 1.00 | 11 | 43 |
|  | 1990 | 0.00 | 0.04 | 0.10 | 1.00 | 0.00 | 0.05 | 0.05 | 1.00 | 11 | 43 |
|  | 1991 | 0.00 | 0.04 | 0.09 | 0.98 | 0.00 | 0.04 | 0.04 | 1.00 | 11 | 43 |
|  | 1992 | 0.00 | 0.04 | 0.09 | 0.99 | 0.00 | 0.05 | 0.05 | 1.00 | 11 | 43 |
|  | 1993 | 0.00 | 0.04 | 0.09 | 0.98 | 0.00 | 0.05 | 0.04 | 1.00 | 11 | 43 |

